# Intestinal Epithelial PTPN2 Limits Pathobiont Colonization by Immune-Directed Antimicrobial Responses

**DOI:** 10.1101/2024.09.24.614848

**Authors:** Pritha Chatterjee, Marianne R. Spalinger, Charly Acevedo, Casey M. Gries, Salomon M. Manz, Vinicius Canale, Alina N. Santos, Ali Shawki, Anica Sayoc-Becerra, Hillmin Lei, Meli’sa S. Crawford, Lars Eckmann, James Borneman, Declan F. McCole

## Abstract

**Background and Aims:** Loss of activity of the inflammatory bowel disease (IBD) susceptibility gene, protein tyrosine phosphatase non-receptor type 2 (*PTPN2*), is associated with altered microbiome composition in both human subjects and mice. Further, expansion of the bacterial pathobiont, adherent-invasive *E. coli* (AIEC), is strongly linked to IBD pathogenesis. The mechanism by which intestinal epithelial cells (IEC) maintain equilibrium between commensal microbiota and immune cells to restrict invading pathobionts is poorly understood. Here, we investigated the role of IEC-specific PTPN2 in regulating AIEC colonization.

**Methods:** Tamoxifen-inducible, intestinal epithelial cell-specific *Ptpn2* knockout mice (*Ptpn2*^ΔIEC^) and control *Ptpn2*^fl/fl^ mice were infected with either non-invasive *E. coli* K12, or fluorescent-tagged *m*AIEC (*m*AIEC^red^) for four consecutive days or administered PBS. Subsequently, bacterial colonization in mouse tissues was quantified. mRNA and protein expression were assayed in intestinal epithelial cells (IECs) or whole tissue lysates by PCR and Western blot. Tissue cytokine expression was determined by ELISA. Intestinal barrier function was determined by *in vivo* administration of 4 kDa FITC-dextran (FD4) or 70kDa Rhodamine-B dextran (RD70) fluorescent probes. Confocal microscopy was used to determine the localization of tight-junction proteins.

**Results:** *Ptpn2*^ΔIEC^ mice exhibited increased *m*AIEC^red^ - but not K12 - bacterial load in the distal colon compared to infected *Ptpn2*^fl/fl^ mice. The higher susceptibility to *m*AIEC^red^ infection was associated with altered levels of antimicrobial peptide (AMPs). Ileal RNA expression of the alpha-defensin AMPs, *Defa5* and *Defa6*, as well as MMP7, was significantly lower in *Ptpn2*^ΔIEC^ vs. *Ptpn2*^fl/fl^ mice, after *m*AIEC^red^ but not K12 infection. Further, we observed increased tight junction-regulated permeability determined by elevated *in vivo* FD4 but not RD70 permeability in *Ptpn2*^ΔIEC^-K12 mice compared to their respective controls. This effect was further exacerbated in *Ptpn2*^ΔIEC^ *m*AIEC-infected mice. Further, *Ptpn2*^ΔIEC^ mice displayed lower IL-22, IL-6, IL-17A cytokine expression post *m*AIEC infection compared to *Ptpn2*^fl/fl^ controls. Recombinant IL-22 reversed the FD4 permeability defect and reduced bacterial burden in *Ptpn2*^ΔIEC^ mice post *m*AIEC challenge.

**Conclusion:** Our findings highlight that intestinal epithelial PTPN2 is crucial for mucosal immunity and gut homeostasis by promoting anti-bacterial defense mechanisms involving coordinated epithelial-immune responses to restrict pathobiont colonization.

## INTRODUCTION

The intestinal epithelium has a strategic position as a physical barrier between trillions of luminal microbes and the immune cells in the underlying lamina propria while also coordinating a very delicate equilibrium to maintain mucosal homeostasis [1–3]. Dysregulation of this physical barrier leads to very serious local and systemic consequences including gastrointestinal infections, inflammatory bowel disease (IBD), type 1 diabetes (T1D), and autoimmune arthritis [4–6].

IBD is a chronic and multifactorial condition affecting more than 6 million people worldwide. [7, 8]. IBD clinically manifests as two sub-types, Crohn’s Disease (CD) and Ulcerative colitis (UC) [9]. IBD involves a complex interplay of host genetics, gut flora, environmental factors, and the immune system. Genome-wide association studies (GWAS) have identified approximately 240 genes associated with IBD [10]. One of the genes identified with disease-associated single nucleotide polymorphisms (SNP) is the protein tyrosine phosphatase non-receptor type 2 (*PTPN2*) gene that encodes the protein, T-cell protein tyrosine phosphatase (TCPTP [11, 12]. Loss of *PTPN2* function leads to aberrant activation/proliferation of T-cells and causes hyper-responsiveness of the Janus Activated Kinase (JAK)- signal transducer and activator of transcription (STAT) pathway [13, 14]. Further, significant alterations in the intestinal microbial community – dysbiosis – have been observed in IBD patients including those carrying *PTPN2* SNPs [15–17]. Our current understanding indicates that genetically pre-disposed hosts have an altered intestinal environment that favors the expansion of commensal microbes with pathogenic potential, “pathobionts”, while reducing the number of protective commensal bacteria. One such pathobiont is adherent-invasive *E. coli* (AIEC), which was first isolated from the ileum of a CD patient [18]. AIEC can adhere and attach to intestinal epithelial cells, in addition to surviving within macrophages [19]. Previously, we have reported that whole body constitutive *Ptpn2*-KO mice exhibit a reduced abundance of *Bacteroidetes* and a greatly increased abundance of *Proteobacteria* compared to *Ptpn2* wild-type littermates. Of note, the greatest increase in abundance within the phylum *Proteobacteria* in *Ptpn2*-KO mice was a novel *Escherichia coli* species that shared significant sequence overlap with human AIEC. We have labeled this novel mouse adherent-invasive *E. coli* (*m*AIEC) strain, UCR-PP2[20].

Intestinal epithelial cells (IECs) are physically connected to each other by tight junction (TJ), adherens junction (AJ) and desmosome protein conglomerates that maintain a tightly regulated semi-permeable barrier while supporting an intact yet flexible epithelial monolayer [21–24]. Disruption of these proteins often leads to increased paracellular barrier permeability [25]. The epithelial monolayer plays an essential role in the gut as it interacts with the commensal population in the luminal space and transmits specific signals that educate the underlying innate and adaptive immune cell population, thereby maintaining mucosal immune homeostasis [26–28]. However, increases in epithelial permeability or damage to the epithelial barrier can permit potentially pathogenic bacteria to interact with the immune cells in the lamina propria, or in more severe cases to access the circulation and cause sepsis. Mucosal immune cells are critical in eliminating invading pathogens as they secrete cytokines and phagocytose bacteria that are essential for pathogen-clearance [29]. Specialized IECs called Paneth cells, are also important for bacterial defenses as they secrete several antimicrobial peptides that kill bacteria or inhibit microbial growth [30–32]. Therefore, the intestinal barrier is essential for mucosal homeostasis in part by restricting the increase in intestinal permeability that is an early and critical event in the development of several inflammatory conditions including IBD [33, 34]. Previously, our lab has demonstrated that *Ptpn2*-KO mice display increased intestinal permeability and reduced antimicrobial peptide production [35, 36]. These results point towards epithelial *Ptpn2* as an important contributor to maintenance of the intestinal barrier and antimicrobial defenses.

However, there remains a major gap in our understanding of how host genetics alters the gut microbial landscape and how opportunistic pathobionts hijack the host defense machinery and manifest complex conditions in a disease susceptible host. Thus, the main aim of this study was to determine the role of *Ptpn2* in the intestinal epithelium in mediating microbiome-immune cell crosstalk to prevent *m*AIEC colonization and preserve gut homeostasis.

## METHODS

### Animal procedures

#### Ethical Statement on Mouse Studies

All animal care and procedures were performed in accordance with institutional guidelines and approved by the University of California, Riverside Institutional Animal Care and Use Committee under Protocol #A2022001B.

### Housing and husbandry of experimental animals

Tamoxifen-inducible IEC-specific *Ptpn2* knockout (*Ptpn2* ^ΔIEC^) were generated as previously described [36]. Cre negative, floxed Ptpn2 (*Ptpn2*^fl/fl^) littermates were used as controls. Five to six-week-old male and female mice were injected with tamoxifen (Sigma-Aldrich, Saint Louis, MO) via intraperitoneal injections at 1 mg/ml in 100μl of corn oil for 5 consecutive days. Twenty-eight-days after the final injection, mice were orally gavaged with 100μl of either PBS, *E. coli* K12 or *m*AIEC^red^ (*m*AIEC strain-UCR PP2) at 10^9^ cfu/ml for 4 consecutive days. Mice were injected intraperitoneally with recombinant IL-22 (Peprotech, Thermo Fisher Scientific, NJ) at 20 ug/ml per mice every 48 hours. Bacterial infections were performed as described above. All mice were 9–12 weeks-old at the time of sacrifice and housed in specific pathogen-free (SPF) conditions at the University of California, Riverside.

### Bacterial infection and immunofluorescence studies

Bacteria from stocks frozen at –80°C in 1:1 vol/vol glycerol: LB were cultured overnight in Luria–Bertani (LB) broth supplemented with chloramphenicol at 37°C, 250–300 rpm, and regrown the next day in fresh LB-chloramphenicol to exponential phase growth. Culture was pelleted, washed with phosphate buffered saline (PBS), and resuspended in PBS. The bacteria used were *m*AIEC^red^, and K12 (a noninvasive *E. coli*, ATCC 25404).

### Tissue RNA isolation and quantitative PCR

Total RNA was extracted from intestinal segments of mice using RNeasy Mini kit (Qiagen, Hilden, Germany). RNA purity and concentration were assessed by absorbance at 260 and 280 nm. One microgram of total RNA was transcribed into cDNA using qScript cDNA SuperMix (Quanta Biosciences, Beverly, MA). Two microliters of 5×-diluted cDNA were amplified using gene-specific primers (Supplementary Table 1) and GoTaq Green, 2× mix. Gene-specific primers were used with the following conditions: initial denaturation 95°C for 5 minutes, followed by 30 cycles 95°C for 30 seconds (denaturation), 55°C for 30 seconds (annealing), and 72°C for 30 seconds (extension). The final extension was 72°C for 5 minutes. Mouse GAPDH was used as an endogenous control.

### *In-vivo* barrier permeability

Mice were gavaged with 80 mg/mL of fluorescein isothiocyanate (FITC)-dextran (4 kDa) (Sigma-Aldrich) and 20 mg/mL of rhodamine B-dextran (70 kDa) (Sigma-Alrich). After 5 hours, blood was collected by retro-orbital bleed into serum collection tubes. Blood was centrifuged at 4°C, 1500 *g*, for 15 min, and serum was analyzed for FITC-dextran and rhodamine B-dextran concentration. Fluorescence of FITC and rhodamine in samples was determined by loading serum into a black plate and measuring excitation wavelengths of 495 nm and 555 nm, and emission wavelengths of 525 nm and 585 nm, respectively, using a SpectraMax iD3 plate reader (Molecular Devices, San Jose, CA), aligned with established protocols [36, 37]. Standard curves for calculating fluorophore concentration in the samples were obtained by diluting the fluorophore stock in sterile MilliQ.

### Isolation of IECs and western blot analysis

Isolated whole intestinal tissues were everted and incubated in Cell Recovery Solution (Corning, New York) on ice for 2 hours, then vigorously shaken by hand to release IECs. IECs were washed twice with ice-cold PBS, then lysed with radioimmunoprecipitation assay (RIPA) buffer (50 mM Tris-HCl pH 7.4, 150 mM NaCl, 1% NP-40, 0.5% sodium deoxycholate, and 0.1% SDS) supplemented with 1× protease inhibitor (Roche, Basel, Switzerland), 2 mM sodium fluoride, 1 mM PMSF, and phosphatase inhibitors for at least 10 minutes on ice. Cells were homogenized on ice using the Q125 Sonicator (Newtown, CT) lysates centrifuged at 16,200*g* at 4°C for 10 minutes, and supernatants collected into new microcentrifuge tubes. Protein concentration was determined using the Pierce BCA Protein Assay Kit (Thermo Fisher Scientific, Waltham, MA). Loading samples were prepared by mixing the same amount of total protein from each sample with Laemmli loading buffer (60 mM Tris-HCl pH 6.8, 2% SDS, 5% β-mercaptoethanol, and 10% glycerol), then boiling the samples at 95°C for 10 minutes. Twenty µg of protein was loaded on polyacrylamide gels, and after separation by gel electrophoresis, transferred onto polyvinylidene difluoride membranes. Nonspecific epitopes were blocked with 5% milk in Tris-buffered saline with 0.1% Tween-20 added for 1 hour at room temperature. Membranes were incubated overnight with primary antibody at 4°C, washed (×3) with Tris-buffered saline with 0.1% Tween-20, and incubated with horseradish-peroxidase–conjugated secondary antibody against the primary antibody species (Supplementary Table 2) for 1 hour at room temperature. Immunoreactive proteins were detected with x-ray films (Labscientific, Inc, Highlands, NJ) using the SuperSignal West Pico PLUS chemiluminescence detection kit (Thermo Fisher Scientific, Waltham, MA).

### Enzyme-Linked Immunosorbent Assay (ELISA)

Colonic whole tissue was excised to perform ELISAs. IL-22 and IL-6 DuoSet enzyme-linked immunosorbent assays were obtained from R&D Systems (Minneapolis, MN) and performed according to the manufacturer’s guidelines.

### Immunofluorescence

Mouse intestinal segments were embedded in OCT and frozen sections cut into 5-μm thick sections. Slides were brought to room temperature, rinsed twice with PBST (PBS +0.1% Tween20), and fixed in methanol (10 min at −20°C). Slides were then incubated in blocking buffer (PBS + 2% donkey serum, 1% BSA, 1% Triton X, 0.05% Tween-20) for 1 hour before overnight incubation with F’ab donkey-anti-mouse antibody (Jackson ImmunoSearch Inc., Jackson, PA). The slides were rinsed with PBST, incubated with primary antibody (diluted in PBS + 5% normal donkey serum) for 30 min at room temperature, rinsed again with PBS before incubation with a biotinylated secondary anti-mouse antibody (Jackson ImmunoSearch Inc.) for 20 min, followed by an additional wash in PBST and subsequent incubation with Alexa Fluor 488-steptavidin, Alexa Fluor 647 secondary antibody (Jackson ImmunoSearch) for 30 min, and finally mounted with ProlongGold Antifade Reagent with DAPI (Thermo Fisher Scientific). Images were acquired with an inverted Zeiss 880 Airy-scan confocal microscope.

### Statistical analyses

Critical significance was set at α = 0.05. Data was analyzed using parametric models. Data are expressed as mean ± SD for *n* number of biological replicates. Between-group inferences were made by using 1-way or 2-way analysis of variance (ANOVA) followed by Tukey’s or Holm-Sidak post-hoc test. All statistical analysis were performed in GraphPad Prism version 10 (GraphPad Inc., La Jolla, CA).

## RESULTS

### Loss of intestinal epithelial PTPN2 promotes mAIEC colonization

Previous studies from our lab showed that constitutive whole-body *Ptpn2*-deficient mice exhibit expansion of a novel *m*AIEC and display reduced expression of antimicrobial peptides [20,35]. To determine the specific role of epithelial *Ptpn2* in limiting *m*AIEC colonization, we used a tamoxifen-inducible *Ptpn2* Villin-cre transgenic mouse model, which either carried the epithelial floxed *Ptpn2* gene (*Ptpn2*^fl/fl^) but was Cre-negative, or the Cre-positive specific deletion of the epithelial *Ptpn2* gene (*Ptpn2*^ΔIEC^). The mice were treated with PBS; a non-pathogenic, non-invasive *E. coli*, K12; or mCherry fluorescent-tagged *m*AIEC^red^ (Figure 1 A). We used *E. coli* K12 as a bacterial control to distinguish specific host responses to bacterial colonization between a non-invasive *E. coli* and the invasive AIEC strain of bacteria. We validated *m*AIEC^red^ adherence and invasion to intestinal epithelial cells (Supplementary 1A, B, D) and its colonization persistence in mice (Supplementary Figure 1 C). *m*AIEC^red^ infected mice displayed a mild decrease in body weight in both *Ptpn2* ^ΔIEC^ and *Ptpn2* ^fl/fl^ mice (Figure 1B), while no changes in colon length were observed (Figure 1C). We also observed differences in colonization of *m*AIEC^red^ in the proximal colon albeit without reaching statistical significance (Figure 1 D). Interestingly, we observed, *Ptpn2* ^ΔIEC^ had higher *m*AIEC^red^ burden in the distal colon tissue compared to *Ptpn2* ^fl/fl^ mice (Figure 1E). Expectedly, *m*AIEC^red^ displayed higher proximal colon colonization in *Ptpn2* ^ΔIEC^ mice compared to its K12 infected counterparts (Figure 1 D, E). Furthermore, we saw no difference in bacterial load of *E.coli* or *m*AIEC in the luminal contents between the 2 genotypes (Supplementary Figure 2 A-D) suggesting that while the luminal bacterial *m*AIEC load remained similar between the genotypes, *Ptpn2*^ΔIEC^ mice were more susceptible to *m*AIEC colonization of IECs. The increased colonization of *Ptpn2*^ΔIEC^ was visually confirmed by immunofluorescence staining. Higher bacterial burden was associated with epithelial cells stained with epithelial cellular adhesion molecule (EpCam) as compared to the macrophages stained with ADGRE1 (F4/80) (Figure 1F). Taken together, these data demonstrate that IEC-specific loss of *Ptpn2* increases host susceptibility to *m*AIEC infection.

**Figure 1.**
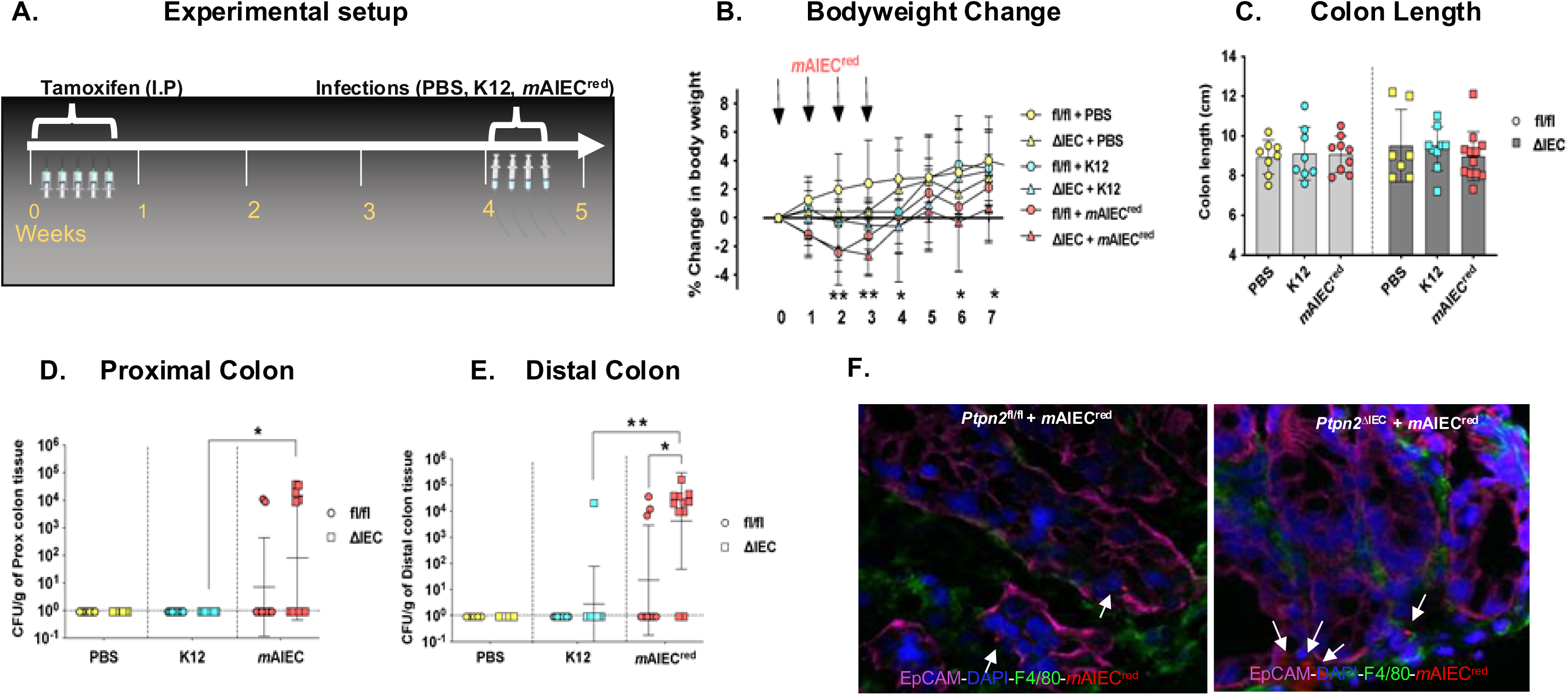
Loss of intestinal epithelial *Ptpn2* promotes *m*AIEC colonization. *Ptpn2*^fl/fl^ or *Ptpn2*^ΔIEC^ were given (A) tamoxifen at 1mg/ml for 5 consecutive days. After a period of 28 days, the mice were given PBS, *E. coli* K12 or *m*AIEC^red^ (10^9^ bacteria per mouse in 100 ul of PBS) from day 0-3 (*n*=9-12). The mice were sacrificed, and bacterial colonies were enumerated at day 7. (B) *m*AIEC^red^ infected groups display mild loss of body weight (*P*=0.006). (C) No appreciable impact was seen in colon length (*P*>0.005). (D) Bacterial colonies were enumerated from proximal colon whole tissues, *Ptpn2*^ΔIEC^ - *m*AIEC^red^ mice have greater *m*AIEC^red^ invasion compared to *Ptpn2*^ΔIEC^ – K12 mice (*P*=0.0194). (E) *Ptpn2*^ΔIEC^ mice display higher *m*AIEC^red^ colonization in the distal colon compared to *Ptpn2*^fl/fl^ controls (*P*=0.002). (F) Immunofluorescence imaging of *Ptpn2*^fl/fl^ or *Ptpn2*^ΔIEC^ proximal colon section infected with *m*AIEC^red^. Epithelial cells were marked in EpCam (magenta), macrophages were marked by F4/80 (green) and *m*AIEC^red^ in red.

### Epithelial PTPN2 restricts bacterial induced intestinal barrier permeability

Previous studies in our lab have shown that *m*AIEC infection exacerbated permeability increases in wildtype C57Bl/6 mice co-challenged with DSS to induce colitis post infection [20]. Therefore, we wanted to determine if AIEC challenged *Ptpn2*^ΔIEC^ can exert an effect in intestinal barrier permeability. We assessed *in vivo* FD4 and RD70 permeability after bacterial challenge. Consistent with our previous data, no significant differences in FD4 permeability were observed between *Ptpn2*^fl/fl^ and *Ptpn2*^ΔIEC^ groups that were treated with PBS (Figure 2A). *E. coli* K12 increased FD4 permeability in *Ptpn2*^ΔIEC^ mice compared to their floxed controls, while *m*AIEC^red^ caused an even greater increase in FD4 permeability in *Ptpn2*^ΔIEC^ versus *Ptpn2*^fl/fl^ mice (Figure 2A). Moreover, no significant difference was observed in RD70 permeability between *Ptpn2*^fl/fl^ and *Ptpn2*^ΔIEC^ mice, or between treatments, suggesting that the epithelial lining was not functionally damaged (Figure 2B). Overall, these data demonstrate that loss of epithelial PTPN2 increases FD4 permeability after bacterial infection, increasing tight junction-regulated paracellular barrier permeability but not non-specific permeability arising from damage to the epithelium.

**Figure 2.**
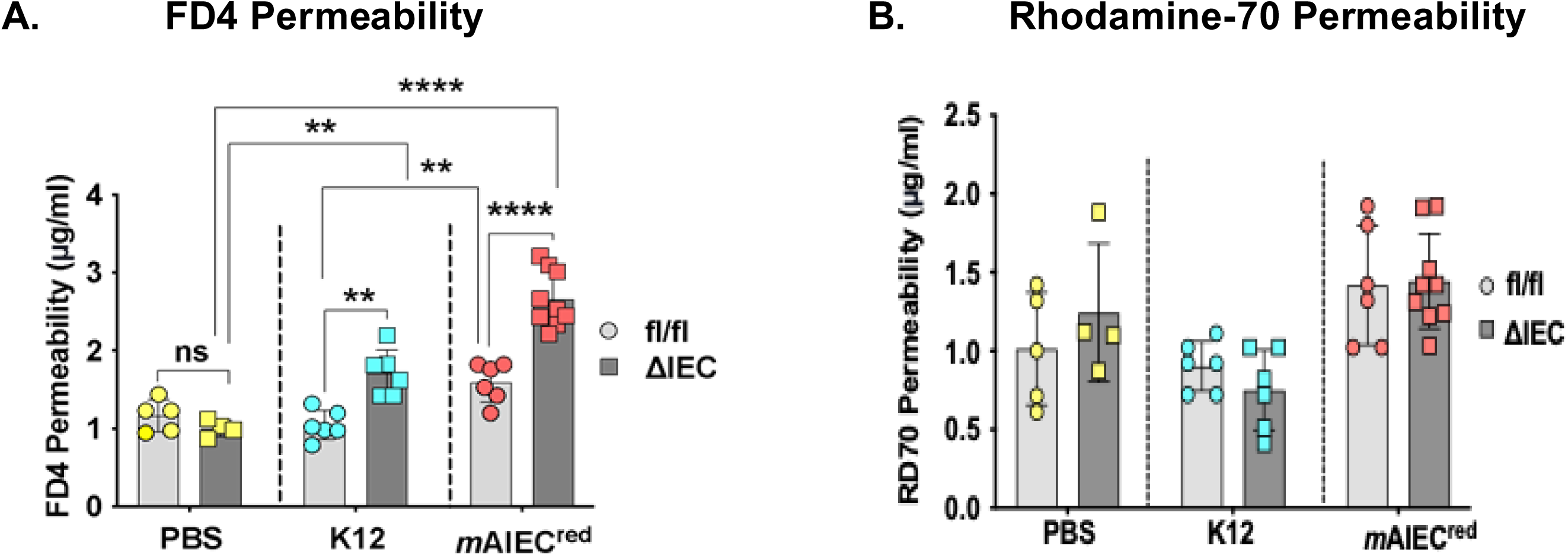
Ptpn2 ^_^ deficient epithelial cells display Increased Barrier Permeability. Intestinal barrier permeability was measured. (A) *Ptpn2*^ΔIEC^ infected with K12 display higher barrier permeability compared to *Ptpn2*^fl/fl^ -K12 controls (*P*= 0.0025), the barrier defect was further exacerbated after *m*AIEC^red^ infection (*P*< 0.0001) in *Ptpn2*^ΔIEC^ compared to floxed controls. The *Ptpn2*^fl/fl^ – *m*AIEC^red^ group also display increased permeability compared to *Ptpn2*^fl/fl^ – K12 mice. (B). No appreciable changes were observed in rhodamine permeability between genotypes or treatment groups.

### AIEC induces greater disruption of barrier-forming proteins in mice lacking PTPN2 in intestinal epithelial cells

Next, we investigated if AIEC-induced changes in intestinal permeability in *Ptpn2*^ΔIEC^ mice were associated with alterations in barrier forming proteins. In line with the FD4 results, proximal colon IECs isolated from *Ptpn2*^ΔIEC^ mice infected with *m*AIEC^red^ had decreased E-cadherin and Occludin protein levels compared to their PBS-treated *Ptpn2*^ΔIEC^ counterparts. (Figure 3 A, B, C, D). Other barrier-forming proteins like JAM-A and Tricellulin remained unchanged between genotypes or treatment groups (Figure 3A). Further, we determined the levels of the claudin family of proteins which are essential for effective barrier function. Claudin-2 is a cation pore-forming member of the claudin family of transmembrane proteins that is frequently elevated during inflammation, including in IBD, while during bacterial infection its expression is increased as part of a host-protective mechanism [39, 40]. Consistent with our previous studies, claudin-2 levels were significantly elevated in *Ptpn2*^ΔIEC^ mice compared to the *Ptpn2*^fl/fl^- PBS treated littermates, while a similar, albeit not statistically significant, effect was observed in the K12 infected *Ptpn2*^ΔIEC^ mice (Figure 3B, E). Strikingly, the increase in claudin-2 levels in *Ptpn2*^ΔIEC^ was reversed after *m*AIEC infection, with these mice displaying a significant reduction in claudin-2. This suggests that *m*AIEC can suppress expression of critical host-protective mechanisms, like claudin-2, even in conditions where claudin-2 is overexpressed (Figure 3B, E). Protein levels of the barrier-enhancing protein, claudin-7, were also reduced in *Ptpn2*^ΔIEC^ – *m*AIEC group compared to respective controls (Figure 3B, F). The Claudin-3 and 4 protein levels remained unchanged by genotype or treatment (Figure 3A, B). Localization of the tight junction regulatory protein, ZO-1, remained intact in all the *Ptpn2*^fl/fl^ groups; however, small gaps were seen in *Ptpn2*^ΔIEC^ – K12 group (Figure 3 G). Furthermore, ZO-1 localization was completely ablated in the *Ptpn2*^ΔIEC^ – *m*AIEC group compared to controls (Figure 3G).

**Figure 3.**
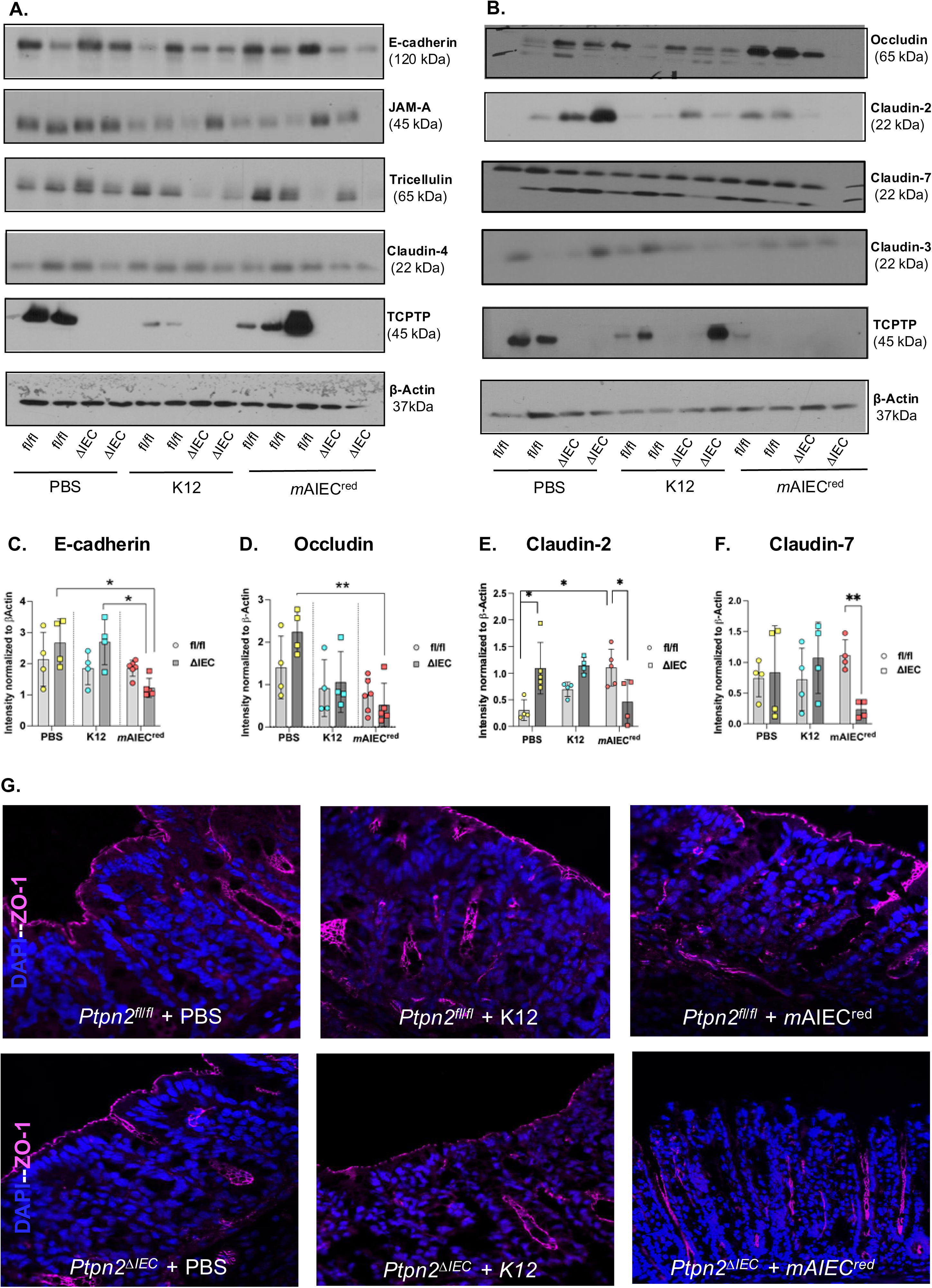
Ptpn2 ^_^ deficient epithelial cells display alterations in barrier-forming proteins post-bacterial infection. (A), (B). Representative western blot images for E-cadherin, occludin, JAMA, tricellulin, claudin-2,3,4 and 7. (C) *Ptpn2*^ΔIEC^ - *m*AIEC^red^ mice show reduced E-cadherin expression compared to *Ptpn2*^ΔIEC^- PBS control (*P*= 0.0124) and compared to *Ptpn2*^ΔIEC^- K12 (*P*= 0.0142). (D) *Ptpn2*^ΔIEC^ - *m*AIEC^red^ ^mice^ display reduced Occludin levels compared to *Ptpn2*^ΔIEC^-PBS controls (*P*= 0.0030). (E) *Ptpn2*^ΔIEC^ display higher Claudin-2 protein expression compared to *Ptpn2*^fl/fl^ controls (*P*= 0.0015). *Ptpn2*^ΔIEC^- *m*AIEC^red^ mice have reduced Claudin-2 levels compared to *Ptpn2*^fl/fl^ -*m*AIEC^red^ controls (*P*= 0.004) whereas *Ptpn2*^fl/fl^ - *m*AIEC^red^ display higher Claudin-2 protein expression compared to their respective PBS controls (*P*= 0.0025). (F) *Ptpn2*^fl/fl^ - *m*AIEC^red^ display lower Claudin-7 protein expression compared to *Ptpn2*^fl/fl^ - *m*AIEC^red^ controls (*P*= 0.0009). Immunofluorescence staining of proximal colon whole tissue. (G) *Ptpn2*^ΔIEC^ mice display gaps in ZO-1 localization (magenta). The gaps in ZO-1 increased in K12 infection in comparsion with the PBS condition and was completely ablated after *m*AIEC^red^ infection.

### PTPN2 deficiency results in reduced antimicrobial peptide production in response to mAIEC infection

Given that *Ptpn2*^ΔIEC^ mice exhibited higher *m*AIEC^red^ bacterial load in distal colon tissue, and our previous observations that whole body constitutive *Ptpn2*-KO mice have reduced antimicrobial peptide (AMP) production and numbers of Paneth cells, we next investigated if the increased susceptibility of *Ptpn2*^ΔIEC^ mice to *m*AIEC^red^ infection involved altered host antimicrobial peptide defenses [35]. Ileal IEC-mRNA expression of the 𝜶-defensins (*Defa5* and *Defa6*), was significantly lower in *Ptpn2*^ΔIEC^ vs. *Ptpn2*^fl/fl^ mice, after *m*AIEC^red^ but not K12 infection (Figure 4A, B). Further, we found ileal IEC protein levels of the AMP, lysozyme, were significantly lower in K12 infected *Ptpn2*^ΔIEC^ compared to its respective control littermates (Figure 4C, E). However, the same effect was not seen in the PBS or *m*AIEC infected groups (Figure 4A, E). No difference was observed in levels of regenerating islet-derived 3 gamma (Reg3ɣ), between genotypes or treatments (Figure 4C, F). In addition, expression of the protease matrix metalloproteinase-7 (MMP-7) - which is responsible for the proteolytic cleavage and activation of 𝜶-defensins – was significantly decreased in the ileum of *Ptpn2*^ΔIEC^ mice post *m*AIEC^red^ infection compared to *Ptpn2*^fl/fl^ littermates (Figure 4C, D). Together, these data demonstrate that intestinal epithelial PTPN2 is critical for AMP-mediated defenses in response to *m*AIEC infection.

**Figure 4.**
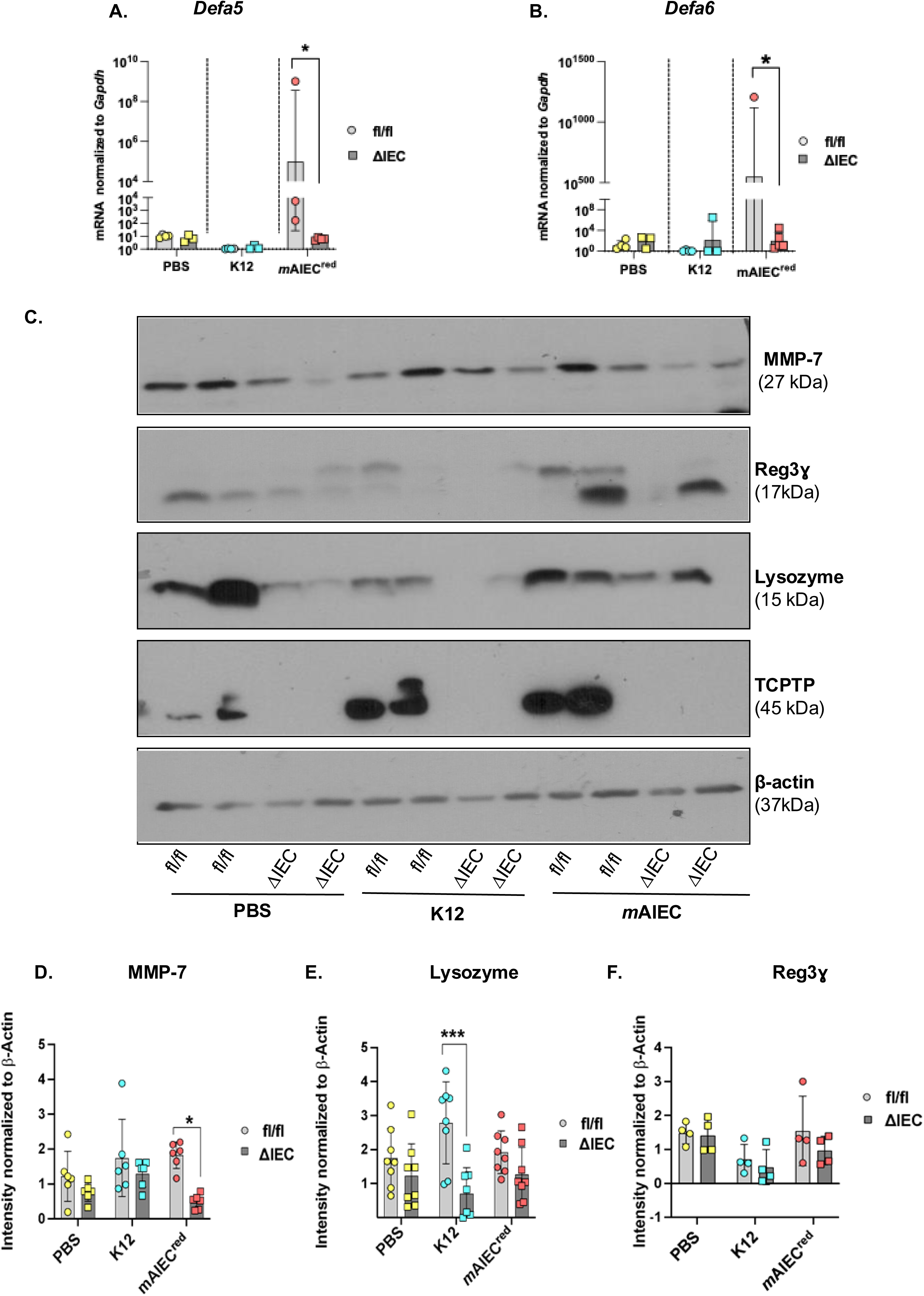
Ptpn2 deficiency results in reduced AMP production in response to mAIEC infection. Antimicrobial peptide expression was measured from ileum whole tissues. *Ptpn2*^ΔIEC^ - *m*AIEC^red^ display reduced (A) *Defa5* (*P*= 0.0008) (B) *Defa6* (*P*= 0.00168) in comparison to *Ptpn2*^fl/fl^ - *m*AIEC^red^. (C) Representative western blot images for MMP-7, Reg3ɣ and Lysozyme from ileum IECs. (D)MMP-7 protein levels were reduced *Ptpn2*^ΔIEC^ - *m*AIEC^red^ compared to *Ptpn2*^fl/fl^ - *m*AIEC^red^ mice (*P*= 0.042). (E) *Ptpn2*^ΔIEC^- K12 mice have lower lysozyme expression compared to *Ptpn2*^fl/fl^ - *m*AIEC^red^ mice (*P*= 0.0165) however, no differences were observed in lysozyme protein levels between *Ptpn2*^ΔIEC^ - *m*AIEC^red^ and *Ptpn2*^fl/fl^ - *m*AIEC^red^ groups. (F) No differences were seen in Reg3ɣ protein levels.

### Intestinal epithelial PTPN2 deficiency caused reduced cytokine expression in response to mAIEC infection

Next, we determined the expression profiles of mucosal cytokines involved in promoting Paneth cell function and clearance of bacteria and bacterial products. Typically, T-helper cells 1 (Th1) mount a cytokine reaction during bacterial invasion which is more inflammatory in nature, whereas the T-helper cells 2 (Th2) mount a response against parasitic infestations. Additionally, mucosal cytokines are important in intestinal barrier maintenance. We observed mRNA expression of *Il22*, *Il6,* and *Il17* was significantly reduced in the proximal colon of *Ptpn2*^ΔIEC^ - *m*AIEC^red^ mice compared to their control littermates (Figure 5A, B, C). Interferon-gamma (*Ifng*) and *1l1b* levels remained similar between genotypes and treatments (Figure 5D, E). Further, western blotting on proximal colonic whole tissues identified reduced expression of the general immune cell marker, cluster of differentiation-45 (CD45), and the T-cell marker, CD3 in *Ptpn2*^ΔIEC^ – *m*AIEC mice in comparison with their respective controls (Supplementary Figure 3 A, B, C). Further, no changes were observed in expression of Type 2 cytokines *Il4*, *Il13* or the Th-2 cell marker, GATA-binding protein 3 marker (*Gata3*) (Supplementary Figure 4A, B, C). Next, we validated the mRNA data by confirming that in colonic whole tissues, PBS treated *Ptpn2*^ΔIEC^ mice had reduced IL-22 and IL-6 protein in comparison to its *Ptpn2*^fl/fl^ control, as determined by ELISA (Figure 5F, G). These data indicate that protective barrier and AMP-promoting cytokine responses are impaired in *Ptpn2*^ΔIEC^ mice.

**Figure 5.**
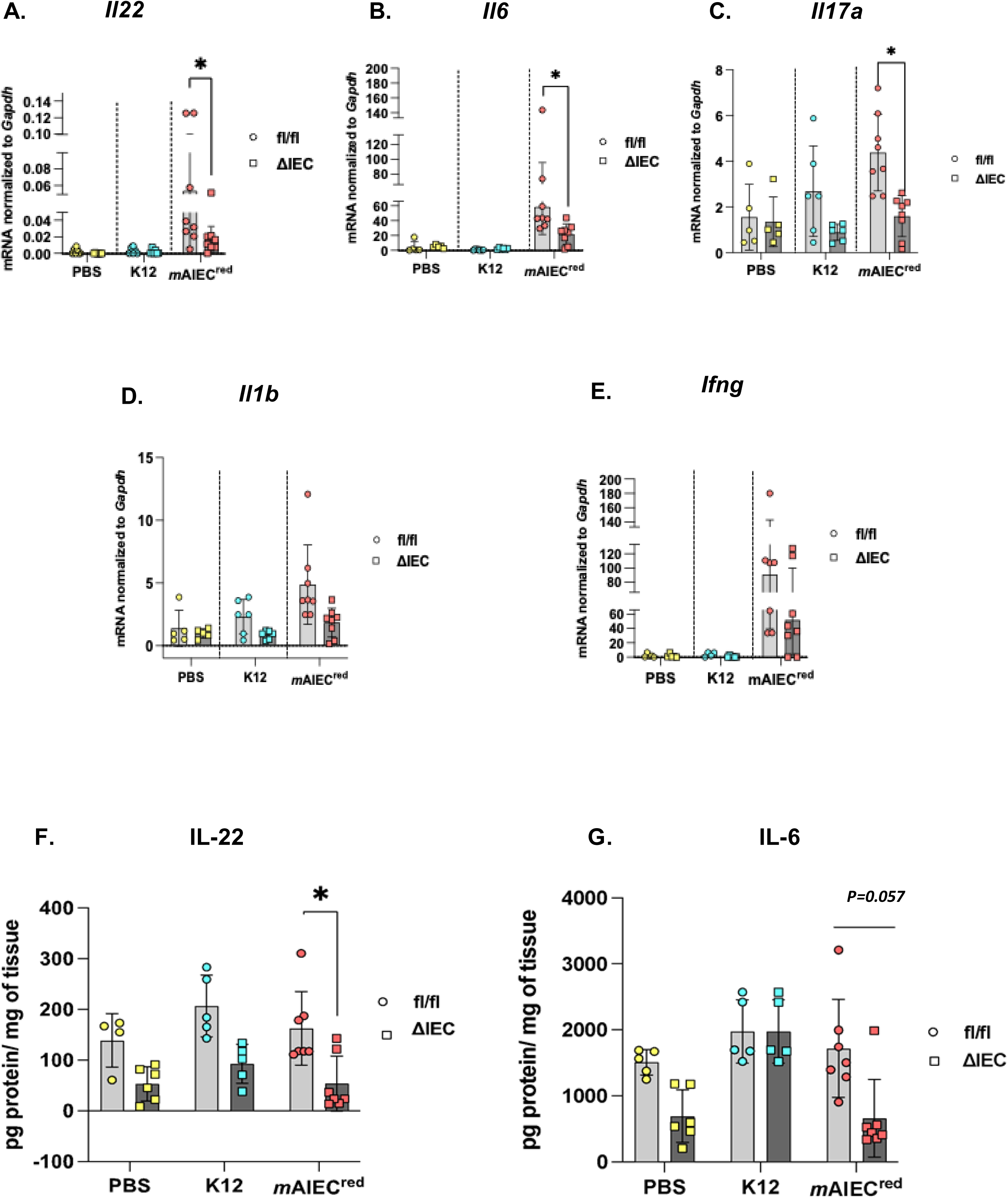
Ptpn2 deficiency results in reduced cytokine expression in response to mAIEC infection. Cytokine expression profile was measured from proximal colon whole tissue sample. (A) *Ptpn2*^ΔIEC^ - *m*AIEC^red^ display lower cytokine levels compared to *Ptpn2*^fl/fl^ - *m*AIEC^red^ for (A) *Il22* (*P*=0.0234) (B) *Il6* (*P*=0.0208) (C) *Il17a* (*P*= 0.00216). No changes were observed between genotype or groups for (D)*Ifng* (E) *Il1b*. ELISA was performed from whole tissue distal colon samples. (F) *Ptpn2*^ΔIEC^ - *m*AIEC^red^ displays lower IL-22 protein levels compared to *Ptpn2*^fl/fl^ - *m*AIEC^red^ (*P*=0.0012). (G) IL-6 levels were also lowered between *Ptpn2*^ΔIEC^ - *m*AIEC^red^ and *Ptpn2*^fl/fl^ - *m*AIEC^red^ groups (*P*=0.005)

### Recombinant IL-22 reduces mAIEC bacterial load and ameliorates the FD4 permeability defect in Ptpn2***^ΔIEC^*** mice in response to mAIEC infection

Given that IL-22 plays a critical role in the production of AMPs and is also essential for the integrity of the mucosal barrier, we next examined if reconstitution of IL-22 in *Ptpn2*^ΔIEC^ mice augmented host defenses to limit *m*AIEC^red^ colonization and restore epithelial barrier function. Vehicle-treated *Ptpn2*^ΔIEC^ mice (+ *m*AIEC) displayed higher *m*AIEC CFU burden in the proximal and the distal colon compared to *Ptpn2*^fl/fl^ mice treated with vehicle alone, whereas IL-22 administration in both *Ptpn2*^fl/fl^ and *Ptpn2*^ΔIEC^ mice significantly reduced the *m*AIEC^red^ CFU burden (Figure 6A, B). Next, we determined FD4 permeability between vehicle and recombinant IL-22 treated *Ptpn2*^fl/fl^ and *Ptpn2*^ΔIEC^ mice post *m*AIEC infection. We also observed that FD4 permeability in IL-22 administered *Ptpn2*^ΔIEC^-*m*AIEC^red^ mice was significantly reduced compared to the vehicle-treated controls (Figure 6C). Interestingly, the same effect was not observed between the vehicle or recombinant IL-22 treated *Ptpn2*^fl/fl^-*m*AIEC group (Figure 6C). Next, we determined the levels of tight junction proteins that were previously diminished post *m*AIEC^red^ challenge. We observed that protein levels of both occludin and E-cadherin were restored after IL-22 administration in *Ptpn2*^ΔIEC^-*m*AIEC^red^ group (Figure 6D). Further, ileal IECs displayed elevated levels of MMP7, and lysozyme in *Ptpn2*^ΔIEC^-*m*AIEC^red^ group after administration of IL-22 (Figure 6E). Additionally, antimicrobial molecules *Defa5* and *Defa6* were also significantly higher in *Ptpn2*^ΔIEC^-*m*AIEC^red^ group reconstituted with IL-22 in comparison with the PBS controls (Figure 6F, G). These data indicate that increased colonization and the subsequent barrier defect post *m*AIEC^red^ infection in *Ptpn2*^ΔIEC^ mice were reversed with IL-22 supplementation.

**Figure 6.**
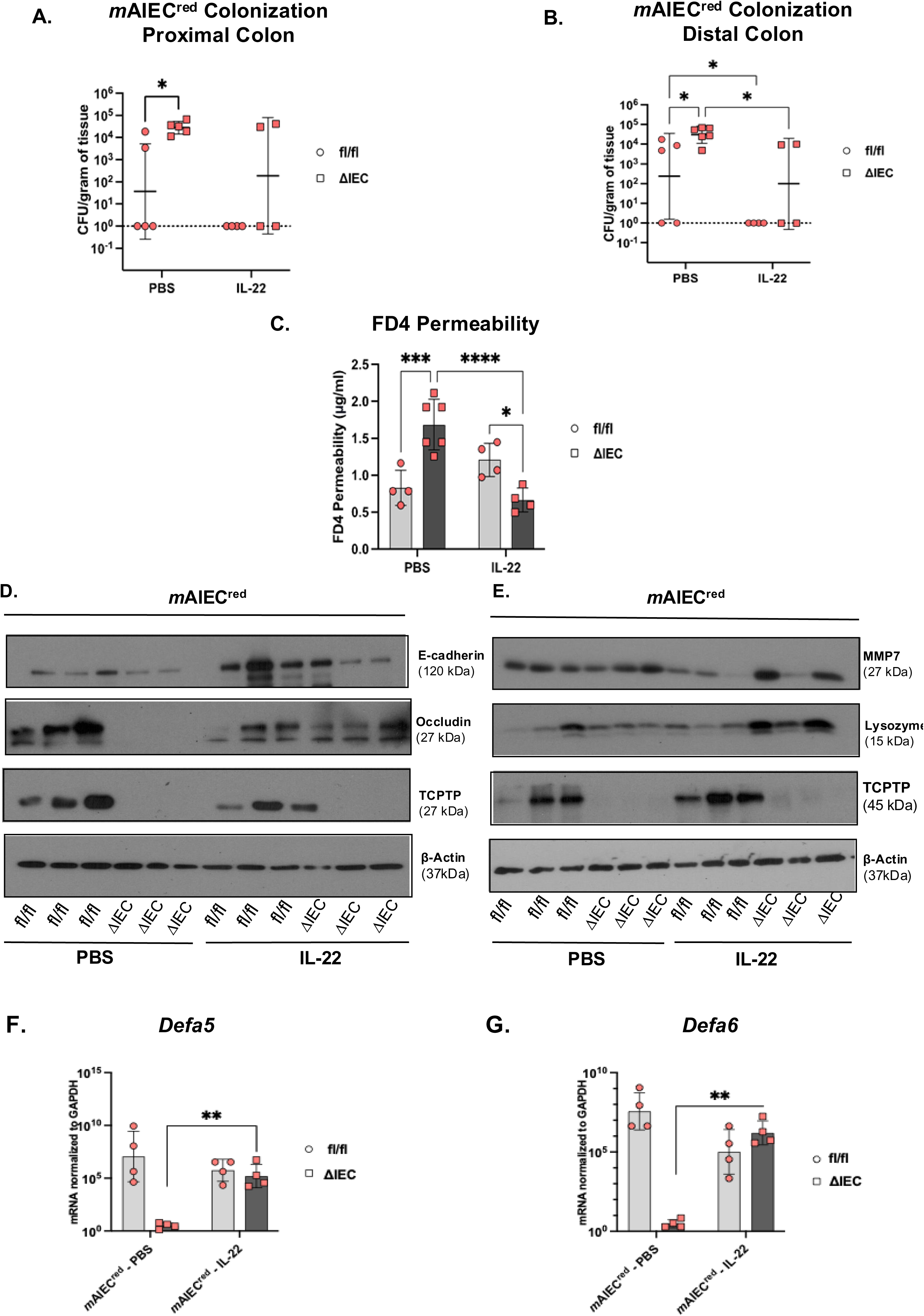
Recombinant IL-22 reduces mAIEC bacterial load and ameliorates FD4 permeability defect in Ptpn2 deficient mice in response to mAIEC infection. Recombinant IL-22 or PBS was administered in *Ptpn2*^fl/fl^ or *Ptpn2*^ΔIEC^ mice starting 2 days prior *m*AIEC^red^ infection and subsequently injected every 48-hours till the mice were sacrificed. PBS injected *Ptpn2*^ΔIEC^- *m*AIEC^red^ mice display higher *m*AIEC^red^ burden than *Ptpn2*^fl/fl^ - *m*AIEC^red^ groups. The *m*AIEC^red^ burden is dramatically reduced in both *Ptpn2*^ΔIEC^ and *Ptpn2*^fl/fl^ IL-22 administered groups in (A) Proximal colon (*P=*0.0168) and (B)Distal colon (*P=*0.0296; *P=*0.0394). (C) FD4 permeability was reduced in IL-22 administered *Ptpn2*^ΔIEC^- *m*AIEC^red^ group in comparison to the vehicle treated *Ptpn2*^ΔIEC^- *m*AIEC^red^ group(*P=*0.0002). (D) E-cadherin and occludin levels in proximal colon IECs were restored in *Ptpn2*^ΔIEC^ - *m*AIEC^red^ mice after administration of IL-22. (E) MMP7 and lysozyme levels were elevated in the ileal IECs of *Ptpn2*^ΔIEC^ -*m*AIEC^red^ after reconstitution of IL-22. (F) *Defa5* and (G) *Defa6* were elevated in after IL-22 administration in *Ptpn2*^ΔIEC^ -*m*AIEC^red^ group compared to PBS treated *Ptpn2*^ΔIEC^ -*m*AIEC^red^ group (*P=*0.0026; *P=*0.00134).

## DISCUSSION

*PTPN2* modulates intestinal microbial composition, antimicrobial peptide levels, and mucosal barrier permeability [20, 35, 36]. We have previously demonstrated that constitutive whole-body loss of *Ptpn2* results in expansion of AIEC and reduction in Firmicutes such as segmented filamentous bacteria (SFB) [20]. Further, we have demonstrated that loss of *Ptpn2* is critical for regulation of intestinal permeability through epithelial-macrophage crosstalk and preservation of barrier-forming proteins [43]. In this study, we demonstrate for the first time that intestinal epithelial PTPN2 (protein TCPTP) is critical for immunity against *m*AIEC infection by promoting antimicrobial defense molecules, maintaining barrier integrity, and coordinating protective immune cell-mediated cytokine responses, highlighting the central role of PTPN2 in microbial-epithelial-immune homeostasis.

We confirmed that our fluorescent AIEC (*m*AIEC^red^) has similar properties of adherence, invasion and survival in macrophages compared to its human clinically relevant counterpart [44]. Further, *m*AIEC^red^ can localize in IECs *in vivo* which validates its identity as an invasive bacterium. While the first AIEC was discovered in ileal tissues of a Crohn’s disease patient, several other AIEC strains have been discovered from other intestinal regions [18, 45–47]. *m*AIEC is present in low levels in wild-type mice but displayed much higher abundance in constitutive *Ptpn2*-KO mice in both the small and large intestine [20]. Of note, in this current study *m*AIEC^red^ preferentially colonized the distal colon versus more proximal regions of the intestine. While the reasoning for this has not yet been determined, it may reflect the impact of specific bacterial niches present in different regions of the intestine, with *m*AIEC achieving the greatest colonization efficiency in the distal colon [20].

Paneth cells are specialized secretory cells present in the small intestine mucosa and produce several AMPs that prevent bacterial colonization and growth. CD patients present with Paneth cell abnormalities with reduced expression of defensins [48]. In this study, we observed that *Ptpn2*^ΔIEC^ mice display reduced expression of the 𝜶-defensins, *Defa5* and *Defa6*, in response to infection with *m*AIEC. Further, these mice also display loss of MMP-7, a protease that is important for activation of the defensin family of proteins, and thus also serves as a Paneth cell marker [49]. Given the critical function of AMPs in restricting bacterial colonization, these data align with the increased *m*AIEC colonization that we observed in the *Ptpn2*^ΔIEC^ mice. Despite the observed defects in Paneth cell antimicrobial peptide expression occurring at their localized site in the small intestine, it is surprising that no bacterial colonization was observed in the ileum (data not shown). It is important to note that Paneth cell AMPs exert important effects not just in the ileum, but their reduced expression can also exert effects more distally and disruption of ileal Paneth cells can provoke dysbiosis in the colon [50]. This may contribute to the region-specific variations in bacterial niches along the lower intestine and the colonization preference of *m*AIEC in the distal colon.

Enhanced intestinal barrier permeability is a risk factor for onset of IBD [51, 52]. Newer evidence has demonstrated that healthy first-degree relatives of patients with CD have significantly higher risk of developing the disease if they are presented with increased intestinal permeability, suggesting that barrier loss is an early and crucial event in disease pathogenesis [33, 34]. Since the intestinal epithelium is semi-permeable, it maintains flux across the epithelial barrier through tight junction-dependent paracellular “pore” and “leak” pathways. The “unrestricted pathway” is generated by epithelial damage due to apoptosis or cell shedding. While the unrestricted pathway – assessed by RD70 – remained unchanged in our study regardless of PTPN2 genotypes or following bacterial challenge, we did observe increased paracellular flux via the leak pathway, as assessed by *in vivo* FD4 challenge. There was no observable difference in *in vivo* FD4 flux between the genotypes in the PBS condition. This was, consistent with our previous reports, although we did find region-specific increases in FD4 permeability in *ex vivo* Ussing chamber studies indicating a mild underlying paracellular flux in *Ptpn2*^ΔIEC^ mice that was exacerbated by cytokine challenge [36]. In our current study, we did make the intriguing finding that FD4 permeability was higher in the *Ptpn2*^ΔIEC^ mice infected with either K12 or *m*AIEC compared to infected *Ptpn2*^fl/fl^ mice. This indicates that loss of PTPN2 in intestinal epithelium changes the dynamics of epithelial interactions with commensal bacteria and appears to confer increased pathobiont potential on both invasive and non-invasive commensals. This is significant as it indicates that in the background of a host genetic defect, “benign” bacteria can provoke defects in intestinal barrier function.

Functional changes in permeability following AIEC infection aligned with altered expression and localization of apical tight junction proteins. We observed that *m*AIEC infection of *Ptpn2*^ΔIEC^ mice decreased levels of the transmembrane tight junction protein, occludin, and the adherens junction protein, E-cadherin compared to *Ptpn2*^ΔIEC^ + PBS controls, however the effect of mAIEC was not significantly different between mouse genotypes. ZO-1 membrane localization was substantially ablated in the *Ptpn2*^ΔIEC^- *m*AIEC mice, while *Ptpn2*^ΔIEC^ – K12 mice displayed gaps in ZO-1 localization. Collectively, these changes in junctional protein expression or localization are consistent with the corresponding increases in FD4 permeability. Furthermore, the pore-forming, tight junction protein, claudin-2, was increased in unchallenged *Ptpn2*^ΔIEC^ mice compared to their *Ptpn2*^fl/fl^ controls. Claudin-2 is upregulated in inflammatory states and is shown to increase paracellular electrolyte permeability via the pore pathway, while it also directly increases paracellular water flux [53, 54]. Interestingly, claudin-2 is also upregulated during infections with *C. rodentium* and it mediates diarrhea and clearance of invasive bacteria, thus reflecting a protective role against infection [40]. Consistent with this study, we observed in *Ptpn2*^fl/fl^ mice that claudin-2 was upregulated in response to *m*AIEC infection. Interestingly, the increased background claudin-2 in *Ptpn2*^ΔIEC^ mice did not mediate a predicted protective effect against infection. Indeed, claudin-2 levels were suppressed in these mice to levels comparable to uninfected *Ptpn2*^fl/fl^ mice. Thus, it appears that loss of PTPN2 in intestinal epithelium compromises multiple host defense mechanisms including AMP production, intestinal permeability, and claudin-2 mediated paracellular water flux.

To understand the mechanisms mediating these effects, we identified that PTPN2 loss compromised local mucosal cytokine responses involved in promoting AMP production (IL-22, IL-17A) and claudin-2 (IL-6, IL-22) expression. IL-22 plays a critical role during pathogen infiltration by stimulating AMPs and preserving the mucosal barrier through epithelial regeneration [40,55]. Further, IL-22 receptors are equally expressed in *Ptpn2*^fl/fl^ and *Ptpn2*^ΔIEC^ mice, therefore, we supplemented *Ptpn2*^fl/fl^ and *Ptpn2*^ΔIEC^ with exogenous IL-22 to determine if the increased susceptibility and barrier dysfunction in *Ptpn2*^ΔIEC^ mice could be rescued by IL-22[41]. We observed that *Ptpn2*^fl/fl^ mice displayed a dramatic reduction of *m*AIEC bacterial load following IL-22 treatment. Furthermore, the bacterial load in *Ptpn2*^ΔIEC^ was also significantly reduced. The FD4 permeability defect observed in *Ptpn2*^ΔIEC^ mice post *m*AIEC^red^ infection was also reversed after reconstitution of IL-22 but the same effect was not seen in the *Ptpn2*^fl/fl^ -*m*AIEC^red^ group. Next, we observed the barrier forming proteins E-cadherin and occludin, which were lower in *Ptpn2*^ΔIEC^-*m*AIEC mice, were restored after administration. Further, we have also demonstrated elevated lysozyme and MMP-7 levels in the *Ptpn2*^ΔIEC^-*m*AIEC group after reconstitution of IL-22. Our results are in line with other studies that showed that IL-22 is necessary for host defense against infiltrating pathobionts [55–57]. *Ptpn2*^ΔIEC^ also displays higher susceptibility to colonization by the pathogen, *C. rodentium*, in comparison to the *Ptpn2*^fl/fl^ mice; however, this phenotype could not be rescued by administration of recombinant IL-22[41]. Rather, the susceptibility was reduced by transferring IL-22 over-expressing macrophages intraperitoneally to the *Ptpn2*^ΔIEC^ mice [41]. These differences likely arise due to differences in host responses to a mucosal pathogen (*C. rodentium*) versus a mucosal pathobiont (*m*AIEC). Taken together these data demonstrate that with the loss of PTPN2 in epithelial cells, a disturbed micro-environment is initiated in the gut rendering it more susceptible to infection or modulation by commensal bacteria with pathobiont potential.

The intestinal epithelium plays a critical yet nuanced role in maintaining homeostasis between the subepithelial immune cells and the luminal microbes. The epithelial monolayer must strike a delicate balance to prevent invading microbes but allow selective permeability necessary for nutrient and electrolyte absorption. Our findings highlight that intestinal epithelial PTPN2 is crucial for mucosal immunity as it promotes antimicrobial peptide defenses and enhances barrier function during infection from pathobionts such as AIEC, and non-invasive commensals. In addition, we identify a unique mechanism for epithelial PTPN2 in maintaining the immune-cytokine landscape of the gut post pathobiont infiltration. These findings reveal an essential role for epithelial expression of this clinically important gene in the maintenance of key defense molecules, barrier function, and the coordinated immune landscape mediating gut homeostasis.

## Supplementary Figure legends

**Supplementary Figure 1.**
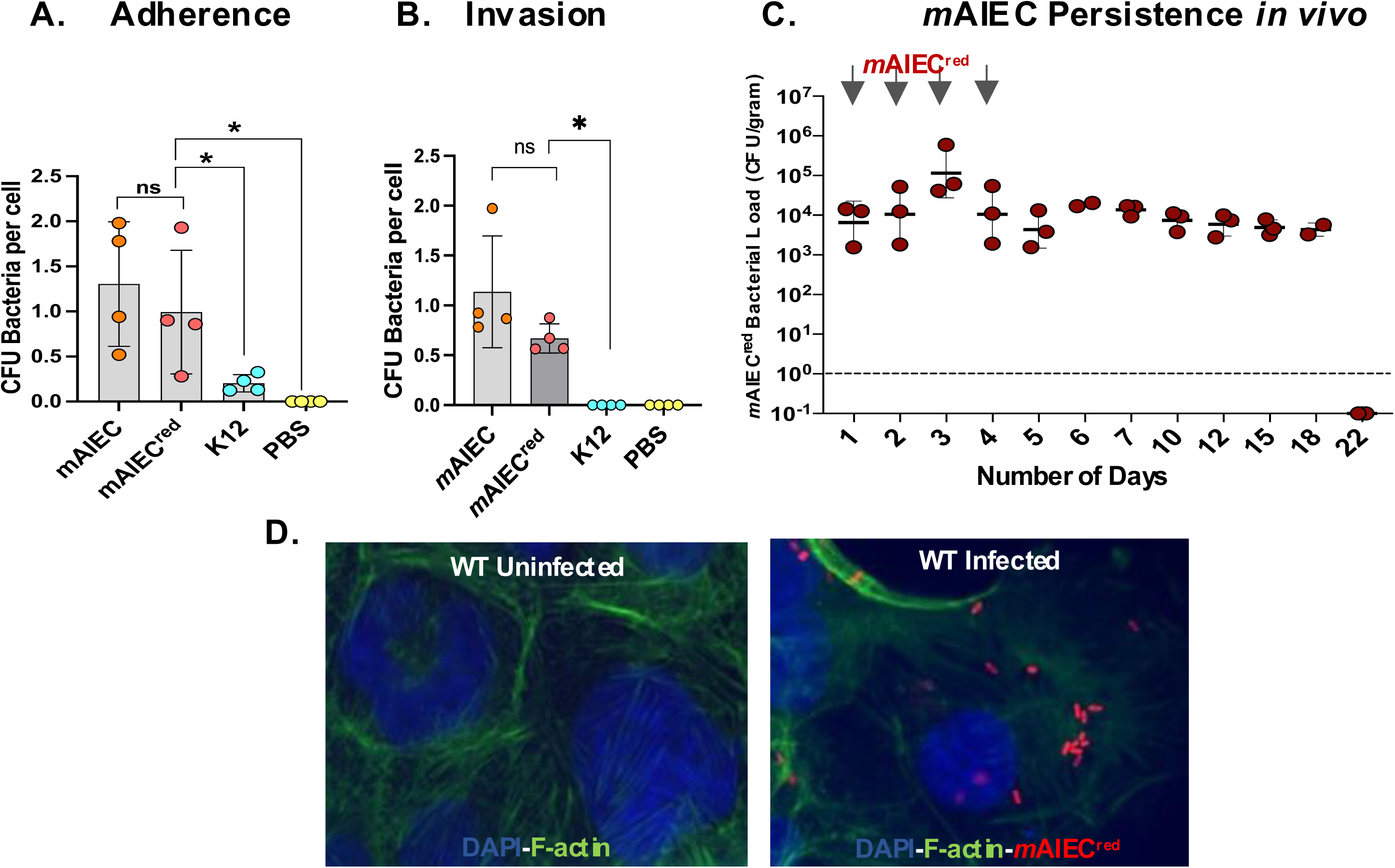
Validation of *m*AIEC^red^. Caco-2BBe cells were infected with PBS, *E. coli* K12, *m*AIEC and *m*AIEC^red^. (A) *m*AIEC and *m*AIEC^red^ display similar (A)Adherence (B) Invasion of Caco-2BBe human IECs. Both had higher invasion compared to *E. coli* K12 (P=0.0023) and PBS was taken as a negative control. (C) C57Bl/6 mice were infected with *m*AIEC^red^ at 10^9^ cfu/ml and was isolated in the mice fecal contents for a period of 21 days. (D) Immunofluorescence was performed on Caco-2BBe cells infected with *m*AIEC^red^. Epithelial cells are marked with F-actin (green) and bacteria is seen in red. *m*AIEC^red^ can invade intestinal epithelial cells.

**Supplementary Figure 2.**
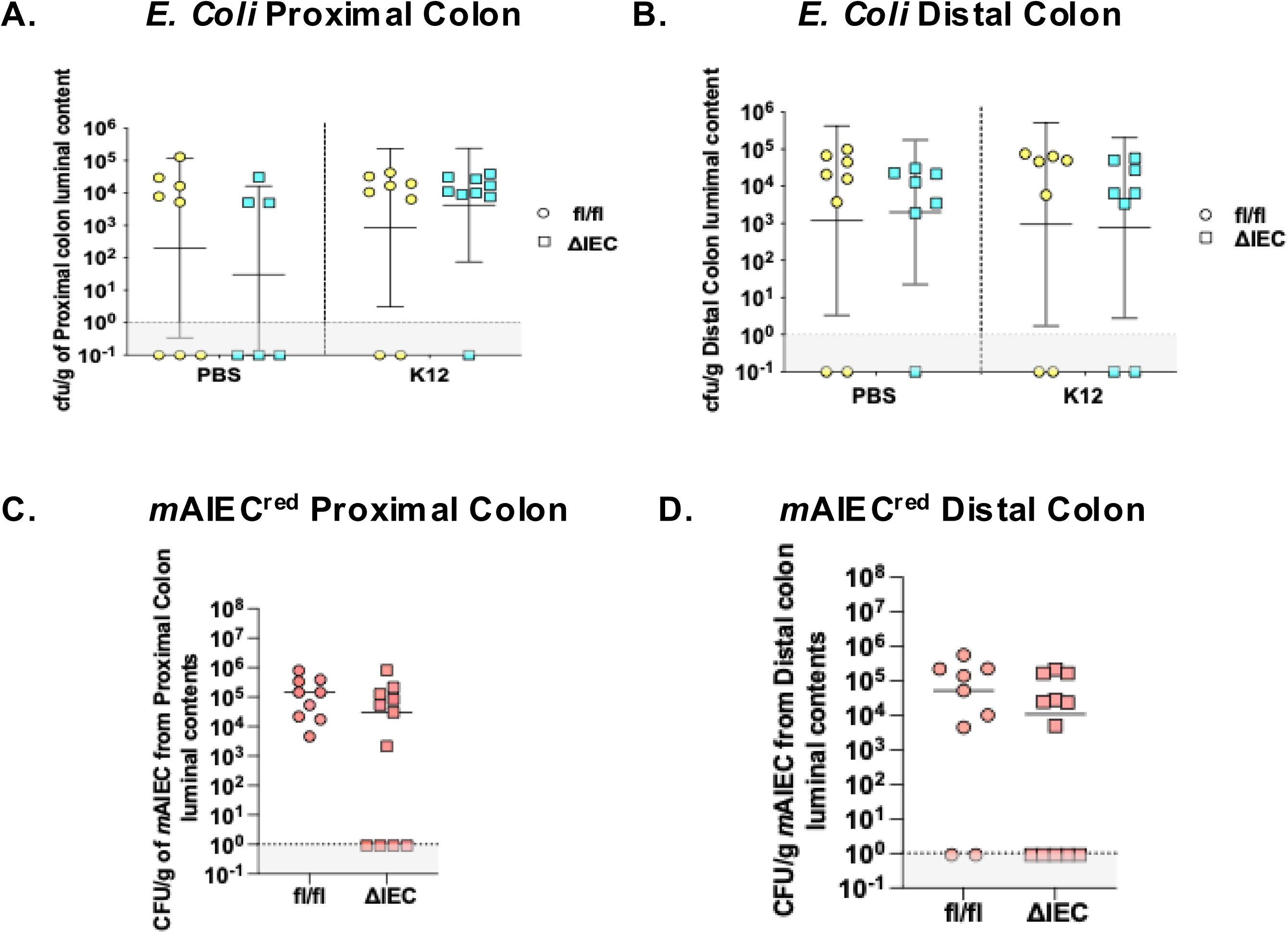
*Ptpn2*^fl/fl^ and *Ptpn2*^ΔIEC^ have similar luminal bacterial load. Bacterial load was enumerated from *Ptpn2*^fl/fl^ and *Ptpn2*^ΔIEC^ mice. (A), (B) Similar *E. coli* burden was observed in the proximal colon, distal colon luminal contents between *Ptpn2*^fl/fl^ and *Ptpn2*^ΔIEC^ mice treated with PBS and K12. (D), E) Comparable *m*AIEC^red^ colonization was observed between *Ptpn2*^fl/fl^ and *Ptpn2*^Δ^ ^IEC^ *m*AIEC infected groups.

**Supplementary Figure 3.**
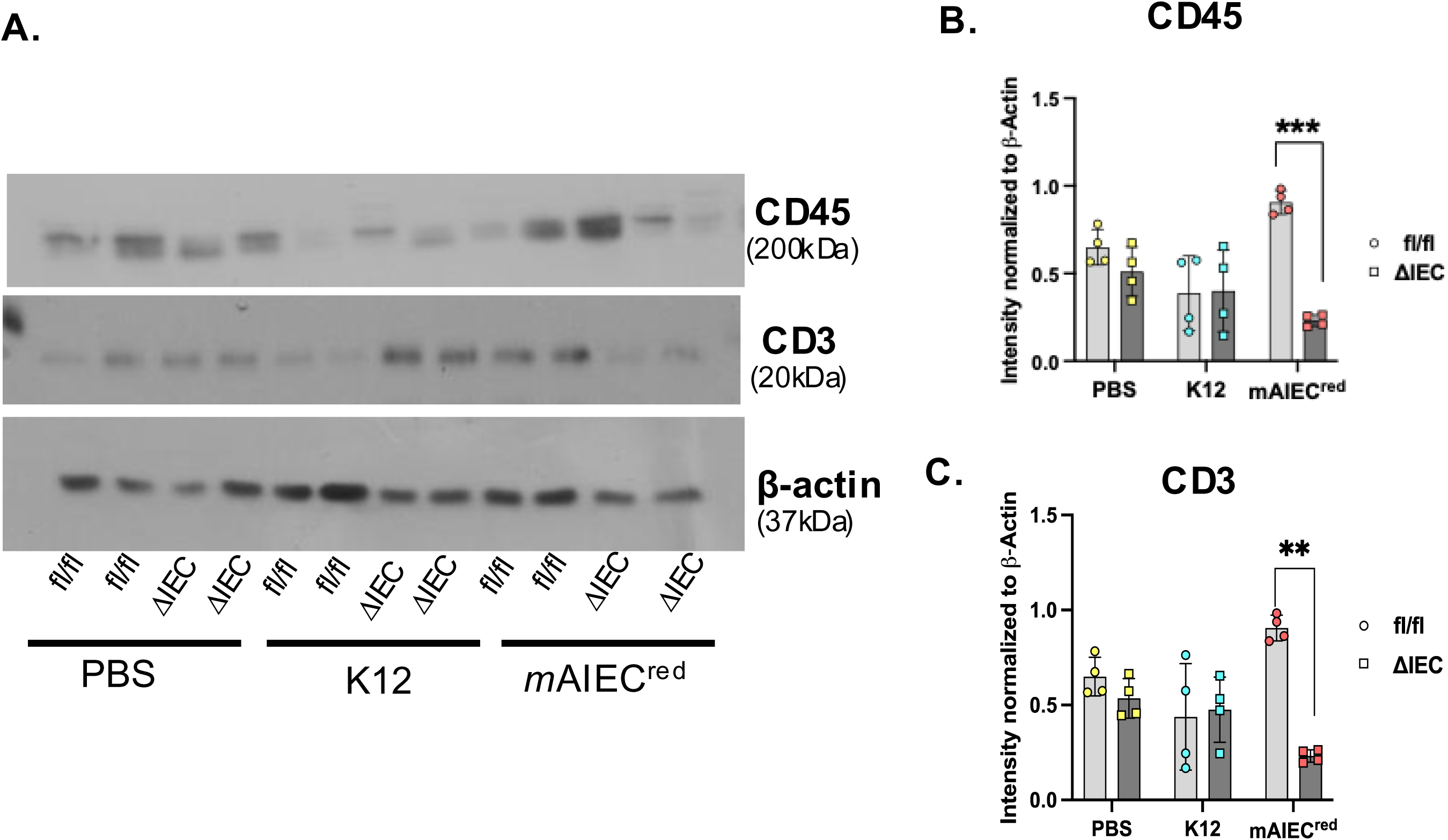
Loss of Epithelial *Ptpn2* led to reduced expression of CD3 and CD45 protein levels. *Ptpn2*^fl/fl^ and *Ptpn2*^ΔIEC^ whole tissue proximal colon was processed for western blot (A), Representative western blot for CD3, CD45 protein expression. *Ptpn2*^ΔIEC^ mice display reduced (B) CD45 protein expression (C) CD3 protein expression after *m*AIEC infection compared to *Ptpn2*^fl/fl^- *m*AIEC group.

**Supplementary Figure 4.**
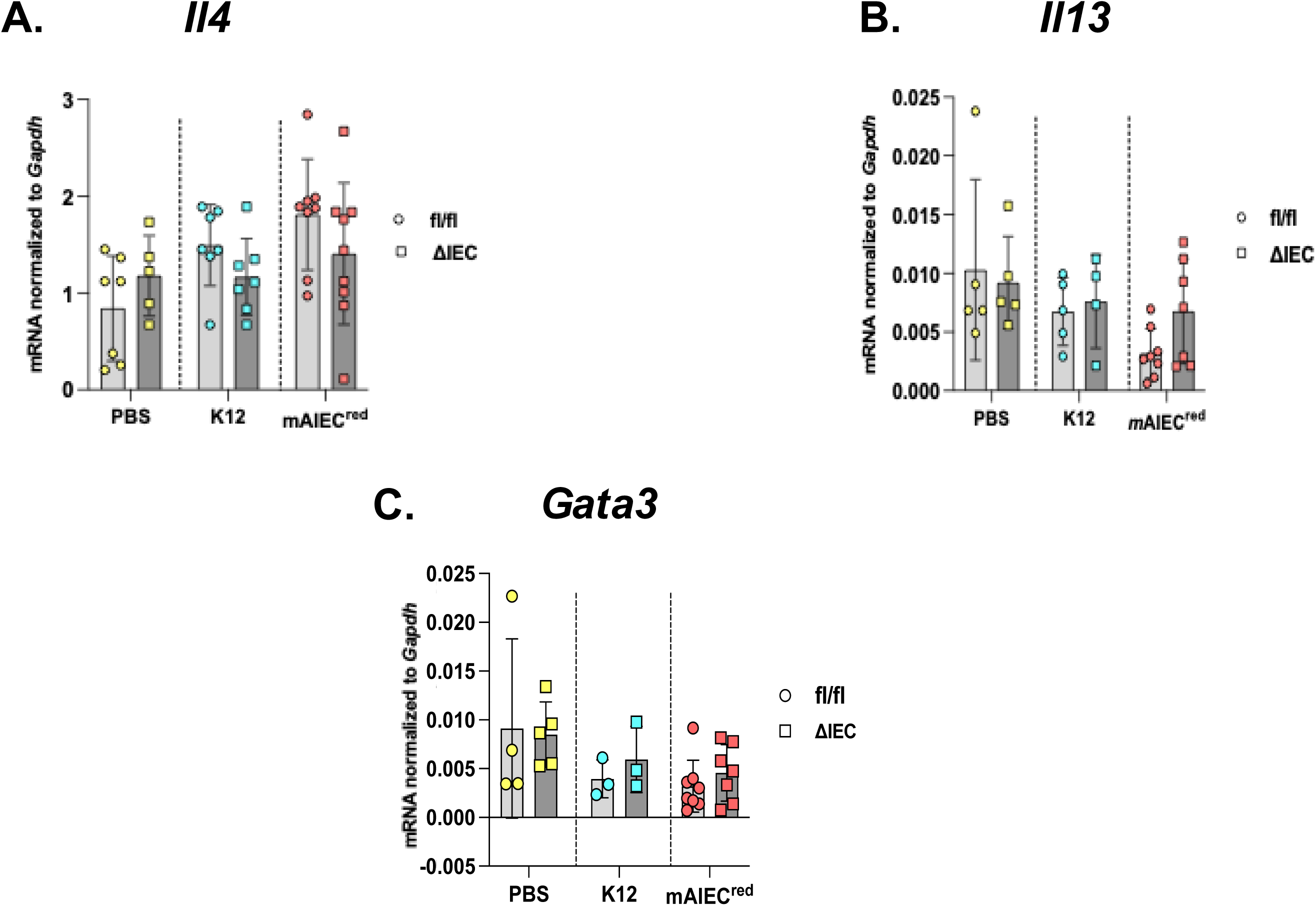
Loss of Epithelial *Ptpn2* does not change Th2 cytokine response. *Ptpn2*^ΔIEC^ and *Ptpn2*^fl/fl^ whole tissue proximal colon samples were processed for mRNA quantification. *Ptpn2*^ΔIEC^ mice do not display any changes in (A) *Il4*, (B) *Il13* or (C) *Gata3* response compared to control or infected littermates.

**Supplementary Table 1:**
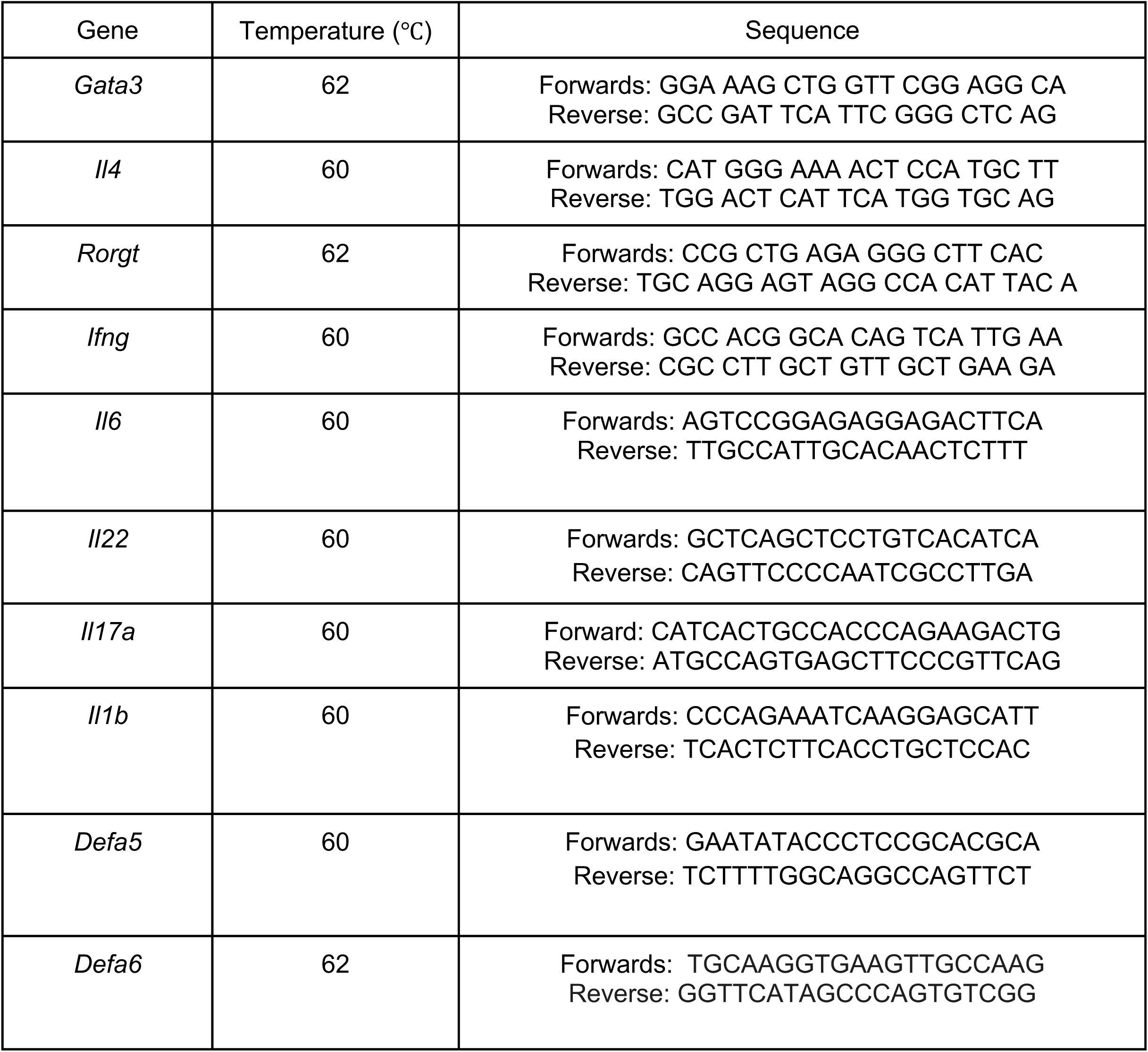
Primer Sequences for Quantitative PCR.

**Supplementary Table 2:** Primary and Secondary Antibodies for Western-Blotting.

| **Antibody** | **Host** | **Provider** | **Catalog Number** | **Dilution** |
| --- | --- | --- | --- | --- |
| CD45 | Rabbit | Cell Signaling | 72787 | 1:1000 |
| CD3 | Rabbit | Abcam | ab5690 | 1:1000 |
| Claudin-2 | Mouse | Invitrogen | 32-5600 | 1:1000 |
| Claudin-3 | Rabbit | Invitrogen | 34-1700 | 1:1000 |
| Claudin 4 | Mouse | Invitrogen | 32-9400 | 1:1000 |
| Claudin-7 | Mouse | Invitrogen | 37-4800 | 1:1000 |
| EpCAM | Rabbit | Cell Signaling | 42515 | 1:1000 |
| E-Cadherin | Mouse | BD Biosciences | 610181 | 1:1000 |
| F4/80 | Rat | Thermo Fisher Scientific | MA5-16624 | 1:1000 |
| JAM-A | Rabbit | Invitrogen | 36-1700 | 1:1000 |
| Lysozyme | Rabbit | Abcam | 76784 | 1:1000 |
| MMP7 | Rabbit | Abcam | 232737 | 1:1000 |
| Occludin | Rabbit | Thermo Fisher Scientific | 71-1500 | 1:1000 |
| TCPTP | Rabbit | Cell Signaling | 58935 | 1:1000 |
| MARVELD2 | Rabbit | Invitrogen |  | 1:1000 |
| ZO-1 Polyclonal | Rabbit | Thermo Fisher Scientific | 61-7300 | 1:1000 |
| β-Actin | Mouse | Sigma-Aldrich | A5316 | 1:1000 |
| HRP-conjugated Anti-rabbit IgG | Goat | Jackson Immunoresearch | 111-036-045 | 1:5000 |
| HRP-conjugated Anti-mouse IgG | Goat | Jackson Immunoresearch | 115-036-062 | 1:5000 |

